# Tissue engineered endothelial keratoplasty with controlled endothelial cell density: proof of concept and paving the way for super TEEKs

**DOI:** 10.1101/2025.05.01.651674

**Authors:** Inès Aouimeur, Louise Parveau, Sofiane Fraine, Zhiguo He, Guillaume Bonnet, Tomy Sagnial, Gauthier Travers, Sédao Xxx, Cyril Mauclair, Anaick Moisan, Philippe Gain, Gilles Thuret, Corantin Maurin

## Abstract

Over the past 20 years, endothelial keratoplasty procedures have revolutionized the treatment of corneal endothelial disorders. These conditions have now become the leading indication for corneal transplantation in Western countries and account for half of all donor cornea usage. Despite their undeniable success, the global shortage of donor tissues and major disparities between nations justify the development of alternatives to donor grafts. Cell therapy using injections of suspended endothelial cells has proven effective, and tissue-engineered endothelial keratoplasty (TEEK) comprising a membrane coated with cultured endothelial cells is under development to better mimic the native endothelial graft. Our team utilizes a femtosecond-laser-cut lens capsule disc as a bioengineering scaffold, taking advantage of this novel tissue’s biocompatibility, transparency, curvature, and availability. In the present study, we provide proof of concept, in 12 TEEKs, that it is possible to control the final endothelial cell density (ECD) by varying the seeding density per mm^2^. Cell characterization was performed through morphometric analysis of the endothelial mosaic stained with anti-NCAM (a lateral membrane marker used as a differentiation marker), using the CellPose artificial intelligence algorithm specifically trained for in vitro endothelium segmentation. Five criteria related to pleomorphism, polymorphism, and elongation were combined into a single endothelial quality score. The median cell viability at 28 days of culture, assessed by Hoechst 33342 and Calcein-AM staining, reached 98% (range: 83–99%). The median viable ECD (number of live cells per surface unit) in the highest-density group was 3,245 cells/mm^2^ (range: 2,778–3,753), paving the way for the bioengineering of supra-physiological TEEKs, or super TEEKs.

**Impact statement:** The process of manufacturing tissue-engineered endothelial keratoplasty (TEEK) allows for the control of endothelial cell density (DCE) and, in particular, the creation of super TEEKs, meaning grafts with supra-physiological DCE that are more likely to better withstand the challenges of surgery and have a prolonged lifespan in recipients.

## Introduction

Diseases of the corneal endothelium are the leading indication for corneal transplantation in Western countries. Their standard curative treatment is endothelial keratoplasty (EK), which consists of removing the central 8 or 9 mm of the pathological endothelium and replacing it with a disc of the same diameter dissected or cut from a donor cornea. Clinical results from numerous case series have long confirmed the extremely favorable risk-benefit ratio of EK: anatomical restoration allowing for the potential of full visual recovery (20/20), long graft survival with a minimally invasive procedure, very low per- and postoperative complication rates, and also a very low rejection rate. Its main drawback remains, to this day, the complete dependence on corneal donation, which is globally insufficient and extremely unequal from one country to another^1^.

With the introduction in Japan of the very first approved corneal endothelial cell therapy^2^, the treatment of corneal endothelial diseases has entered a new era: that of Advanced Therapy Medicinal Products (ATMPs). The first product developed is a suspension of human corneal endothelial cells (hCEC) cultured in vitro from a small number of donor corneas under the age of 30. The goal of these innovative treatments is to replace donor corneas in these indications.

In addition to the injection of suspended cells, bioengineered corneal endothelial grafts could represent a promising therapeutic alternative. Tissue Engineered Endothelial Keratoplasty (TEEK) precisely replicates the endothelial graft from a donor: it is composed of a transparent membrane, equivalent to Descemet’s membrane (a basal membrane equivalent for hCEC), covered with a single layer of hCEC to form a functional neo-endothelium.

We have developed a TEEK using the human anterior lens capsule as a support. An original femtosecond laser cutting process allows us to reproducibly obtain an 8-millimeter diameter lens capsule disc (LCD), regardless of the lens diameter, which varies in humans from 9 to 11 mm, mainly depending on age. Its efficacy has been demonstrated in rabbits, where a xeno-TEEK (LCD and hCEC) restored corneal transparency in a model of complete endothelial injury^3^.

The relatively simple manufacturing process allows for control of the number of hCEC seeded on the LCD, then for measurement of the endothelial cell density (ECD in cells/mm^2^) and assessment of cell quality at the end of culture, prior to delivery. Several questions remain unresolved: does this manufacturing process allow for the production of TEEK with a predetermined ECD, and in particular, TEEK with a very high, supra-physiological ECD, which could allow for prolonged graft survival? In this study, we seeded LCDs with three increasing ECD levels and compared the number of viable hCEC as well as cell morphology after 28 days of culture.

## Method

### Human corneas and crystalline lens collection

Crystalline lenses were collected during corneal retrieval (PFS16-010) (**Table S1**) Special care was taken to preserve the lens capsule by avoiding direct pinching. Lenses donors’ age was 73 ± 9 [56 ; 90] (mean ± standard deviation [min ; max]). Lenses were stored in 20 mL of organ culture medium (CorneaMax, Eurobio, Les Ulis, France) at 31°C for up to 62 days (20 ± 13 [1 ; 62]), to evaluate the feasibility of long-term storage before processing.

Primary hCEC cultures were established using corneas retrieved for research, as authorized by the French Biomedicine Agency (PFS16-010 and PFS21-001). Corneas were stored in organ culture medium (CorneaMax) at 31°C prior to hCEC isolation. Corneas from three donors aged 15, 83, and 88 years were used. The death-to-harvest time was less than 24 hours. All samples were handled in accordance with the tenets of the Declaration of Helsinki.

### Preparation of the anterior lens capsule disc

Lenses were placed in a sterile watertight Petri dish, a custom holder machined in polyetheretherketone (PEEK) to allow accurate centration during laser cutting. The anterior capsule was cut using a femtosecond laser prototype system. The source was a Satsuma HP (Amplitude, Bordeaux, France) emitting at 1030 nm, with 328 fs pulse duration. Laser pulses passed through a Galvanometer scanner (Scanlab, Puchheim, Germany) and were focused with an f-theta lens. The focus was a Gaussian-shaped 8 μm diameter spot (1/e^2^), spaced 5 μm apart. Pulse energy was 4 μJ. The capsule was cut to produce an anterior lens capsule disc (LCD) of 8 mm in diameter with 3 peripheral asymmetric marks (2 aligned, 1 offset). This layout, originally used in endothelial keratoplasty, ensured proper tissue orientation. A circular x-y scan was applied at 250 kHz, with 5 μm spacing along the circle edge, as shown in Figure 1. This was repeated vertically (z-axis) in 2 μm steps over 4.6 mm centered on the capsule. The high-density tubular pattern ensured full cutting of the capsule. LCDs were then dissected using microsurgical tools under a binocular microscope (SZ61, Olympus, Tokyo, Japan). After staining with 0.06% trypan blue (VisionBlue, DORC, Zuidland, The Netherlands) to facilitate visualization, the LCD was detached from the lens using toothless forceps. LCD were spread on a glass slide with sterile water to verify their integrity under a microscope.

**Figure 1.**
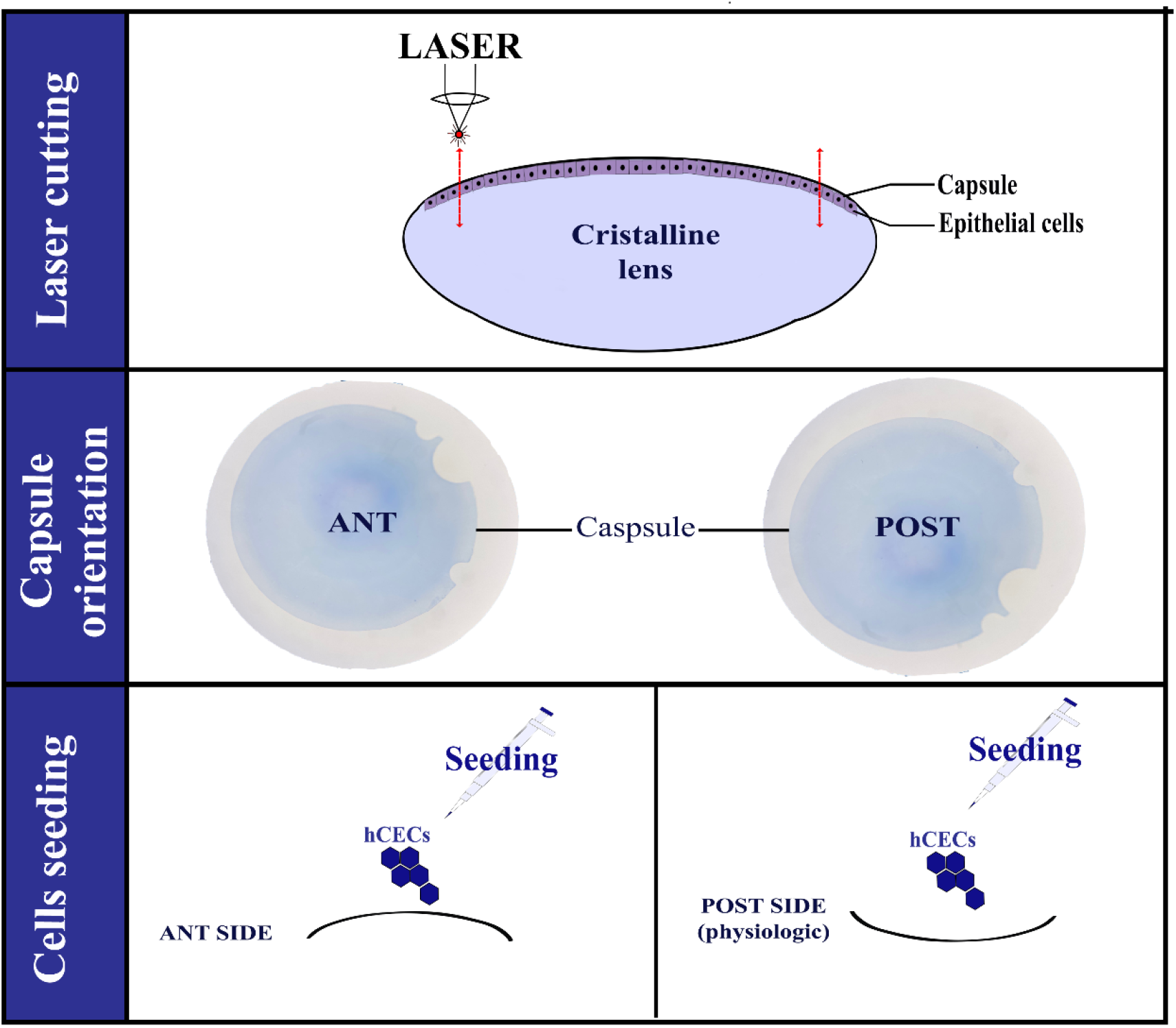
Principles of cutting anterior lens capsule discs (LDC), orientation, and seeding of corneal endothelial cells on the anterior or posterior surface.

They were placed in a glass vial (11553542, Fisher Scientific, Loughborough, UK) containing 4 mL of sterile water (B230531, Versylen, Fresenius, Bad Hamburg, Germany) and 1/100 antibiotic-antimycotic (15240-062, Gibco, Saint Louis, MO, USA). These tubes were then agitated at room temperature for 3 days to ensure the decellularization of the lens epithelial cells. The LCDs were then stored at 4°C in sterile water until use.

### Cell culture

All cell cultures were performed using the “peel and digest” method previously described^4^ and used for the first clinical applications of endothelial cell therapy by injection^2^. The composition of the three culture media was detailed (Table 1).

**Table 1.** Composition of the three media used in the corneal endothelial cell culture process. The passage medium was the expansion medium supplemented with Y-27632.

| Use | Component designation | Company | Reference | Final concentration |
| --- | --- | --- | --- | --- |
| Digestion | Optimem-I | Gibco | 11058-021 |  |
|  | CaCl<sub>2</sub> | Sigma | C5670 | 200 µg/mL |
|  | Ascorbic acid | Sigma | A5960 | 20 µg/mL |
|  | Collagenase A | Gibco | A12177 | 2 mg/mL |
| Expansion | Optimem-1 | Gibco | 11058-021 |  |
|  | CaCl<sub>2</sub> | Sigma | C5670 | 200 µg/mL |
|  | Ascorbic acid | Sigma | A5960 | 20 µg/mL |
|  | Foetal calf serum | Eurobio | CVFSVF00-01 | 8% |
|  | Gentamycine | Gibco | 15710,049 | 50 µl/mL |
|  | Chondroïtine Sulfate 2% | Wako | 0341462 | 0.8 mg/mL |
|  | Epidermal Growth factor h (EGFh) | Gibco | PHG0311 | 5 ng/mL |
|  | SB203580.<br>Adezmapimod. p38 MAP<br>kinase inhibitor | Selleckchem | S107612 | 1,1513µg/mL (10µM) |
|  | SB 431542. TGF-beta<br>inhibitor | Selleckchem | S1067 | 0,1151µg/mL (1µM) |
| Passage | Milieu d'expansion |  |  |  |
|  | +Y-27632. Rock Inhibitor | Selleckchem | S1049 | 10µM |

Each cornea was treated independently. Briefly, the Descemet’s membrane (DM) was peeled off using forceps, under an operating microscope (Universal S3, ZEISS, Germany), using a “no-touch” technique to obtain the largest possible surface of endothelium without trabeculum. The DM was then exposed to 0.06% Trypan blue (Visionblue, DORC, Zuidland, The Netherlands) for 30 seconds to reassess endothelial quality (absence of large areas of dead cells). It was then quickly rinsed in CorneaMax and incubated in a well of a 24-well plate (3524, Costar, Corning Incorporated, USA) containing 500 µL of digestion medium for 16 hours in a humidified incubator at 37°C and 5% CO2. 2.5 mL of passage medium was then added to stop the enzymatic reaction of collagenase A, and the contents of the well were transferred to a conical tube (64199, Caplugs Evergreen, Dutsher). After centrifugation (D37520, Sigma, Germany) at 200G for 5 minutes, the supernatant was removed, and the cell pellet was resuspended in 100 µL of passage medium. The dissociation of the cells was completed by performing about ten successive pipetting steps. The 100 µL were then seeded into a well of a 12-well plate with 380 mm^2^ surface area (3513, Costar, Corning Incorporated, USA) in 1.5 mL of passage medium. The passage medium was replaced after 24 hours with expansion medium. The total volume of each well was then fully renewed once a week with expansion medium.

To perform the passages, the cells were rinsed with Dulbecco’s phosphate-buffered saline (PBS) without calcium or magnesium (LM-S241, Biosera, France), incubated at 37°C for 15 to 30 minutes in a 5X TriplE solution (A12177-01, TriplE 10X, Gibco, Life Technologies Limited, UK), and then mechanically detached by performing 5 to 6 successive pipetting steps. The suspension was centrifuged in a conical tube at 200G for 5 minutes with 2.5 mL of passage medium. The cell pellet was resuspended in 100 µL of passage medium by pipetting several times to dissociate the cells. Passages were all performed at a surface ratio of 1:2 for 24-well to 12-well plates and 12-well to 6-well plates, and at a ratio of 1:2.5 for 6-well plates, in order to double the culture surface at each passage.

All wells, except for the initial digestion step, were coated with a laminin 511 solution (892012, iMatrix Nippi, Incorporated, Tokyo, Japan) diluted in PBS (without calcium or magnesium) at 0.25 µg/cm^2^ and left in place as recommended by the supplier for one hour at 37°C. The entire coating solution was then replaced with passage medium (volume varying depending on the chosen well plate) and placed in the incubator until seeded within the next 20 to 30 minutes. No drying time was performed during the coating step, in accordance with the supplier’s instructions.

### Assembly of Tissue Engineered Endothelial Keratoplasty (TEEK)

The preparation method for the LCDs was the same as recently described^5^. The LCDs were rinsed 3 times in sterile PBS before being spread, under a microbiological safety cabinet (MSC), on the polyethyleneterephthalate (PET) membrane of 113 mm^2^ inserts (665610, Greiner BIO-ONE) that were previously coated with 150 µL of FNC Coating Mix (a solution of fibronectin, collagen, and albumin 0407, Athena ES, Baltimore, MA, USA). To test the influence of the LCD orientation on the growth of hCEC, half of the LCDs were spread in one direction and the other half in the opposite direction (Figure 1). After 2 hours of drying under MSC, the capsules were coated with iMatrix (0.25 µg/cm^2^) for 1 hour in the incubator at 37°C.

The hCEC from the 3 donors (**Table .2**) with a characteristic endothelial phenotype (joint hexagons without mesenchymal endothelial transition) were selected. Three groups of TEEK were generated by varying the number of seeded cells between 500, 2500, and 4000 cells/mm^2^. The hCEC were harvested in the same manner as during a standard passage. The cell concentration per µL was determined (in duplicate) using an automatic counter (TC20, BioRAD, Hercules, CA, USA), and the volume was adjusted to obtain the desired cell density. For each group, the hCEC were seeded on both the anterior and posterior sides of the LCD to analyze the importance of this polarity. The TEEK were then incubated at 37°C and 5% CO2 for 28 days, following the same expansion medium renewal schedule as for primary hCECs: the first renewal 24 hours after seeding, then once a week with 1mL in the bottom of the wells and 600µL in the inserts.

**Table 2.** Characteristics of the corneas used for primary cultures of endothelial cells. ECD before use, corresponded to the ECD of the primary culture before cell harvesting and seeding on the lens capsule discs. Culture assignment corresponded to the number of TEEKs performed using each culture.

| Sex/Age<br>(years) | Death-to-<br>retrieval<br>time<br>(hours) | Initial<br>corneal<br>ECD<br>(cells/mm<sup>2</sup>) | *In vitro*<br>ECD before<br>use<br>(cells/mm<sup>2</sup>) | Passage | Culture<br>assignment |  |  |
| --- | --- | --- | --- | --- | --- | --- | --- |
|  |  |  |  |  | TEEK<br>500 | TEEK<br>2 500 | TEEK<br>4 000 |
| M 15 | 18 | 2 809 | 1 865 | P9 | 2 | 2 | 2 |
| F 83 | 16 | 2 114 | 1 500 | P4 | 2 | 2 | 2 |
| F 88 | 22 | 2 337 | 1 600 | P4 | 1 | 1 | 1 |

M: male, F: female, ECD: endothelial cell density

### Final quality controls

After 28 days of storage, two parameters were measured: 1/ the viable endothelial cell density (number of viable hCECs per unit area or vECD) and 2/ the morphometry of cells labeled with an anti-NCAM used as a reference marker for well-differentiated CECs, without EndoMT ^6^,^7^

The measurement of vECD was obtained after double staining with Hoechst 33342/Ethidium homodimer, using a protocol adapted from Pipparelli et al. ^8^. Briefly, the TEEKs adherent to the membrane of the culture inserts were rinsed with 500 µL of PBS then incubated for 45 minutes at room temperature (RT), protected from light, with a solution of Hoechst at 5 µg/mL (B2261, Sigma, Saint Quentin Fallavier, France) and Calcein-AM at 4 µM (FP-FI9820, Interchim, Montluçon, France) diluted in Opti-MEM I. The TEEKs were then observed directly in the inserts under a macroscope (macrozoom microscope, MVX10, OLYMPUS, Tokyo, Japan) at x1 and x6.3 magnification (Figure S1). The surface covered by viable cells (Calcein-positive) was measured using the Cornea J plugin on FIJI^9^ on x1 macroscope images.

The 12 TEEKs still adherent to the bottom of the inserts (n=3 TEEK 500; n=5 TEEK 2,500; and n=4 TEEK 4,000; 6 seeded on the anterior side and 6 on the posterior side) were then fixed in methanol (A456-212, Fisher Scientific, Loughborough, UK) for 45 minutes at room temperature (RT). The immunostaining protocol was similar to the already published protocol^10^. Briefly, the inserts were rinsed with 500 µL of PBS, then 100 µL of blocking solution, composed of 2% goat serum (11475055, Fisher Scientific) and 2% BSA (A3059, Sigma Aldrich) diluted in PBS, were added into the inserts. The plate with the inserts containing the blocking solution was incubated under gentle agitation at 37°C. After 30 minutes, the blocking solution was replaced with the primary antibody solution composed of the blocking solution supplemented with hNCAM.I (IgG2b, R&D, MAB24081, R&D Systems, Minneapolis, USA) diluted at 1/500. After one hour at 37°C under agitation and protected from light, the compartments were gently rinsed three times with 500 µL of PBS. Finally, the secondary antibody Goat anti-mouse IgG2b AlexaFluor 555 (A110, Invitrogen) diluted at 1/1000 in the blocking solution and 4’,6-diamidino-2-phenylindole dihydrochloride (DAPI) at 1 mg/mL (D9542, MilliporeSigma, St Louis, MO, USA) were incubated for one hour at 37°C under gentle agitation and protected from light. The insert membrane carrying the TEEK was then carefully cut and mounted flat between slide and coverslip in a drop of fluorescent mounting medium (Neo-Biotech, MB23.00158.1, Nanterre, France). The TEEKs were observed under an epifluorescence microscope (IX81, Olympus) with CellSens software (Soft Imaging System GmbH, Munster, Germany). Observations were made with the x4 objective and the multi-image alignment mode to cover the entire surface of the TEEK, then with the x10 and x40 objectives in different areas of interest of the TEEK (Figure S2). The number of nuclei per unit area was obtained on Fiji using the macro function deep learning-based nuclei/cell detection and segmentation method (Stardist) from images acquired under the epifluorescence microscope at x10 with a resolution of 2048×2048 pixels (Figure 2). For the calculation of the vECD, the average value from the center and intermediate zone (weighted by the number of nuclei counted) was used: vECD = %Calcein-positive surface × (nuclei/mm^2^ in these 2 zones).

**Figure 2.**
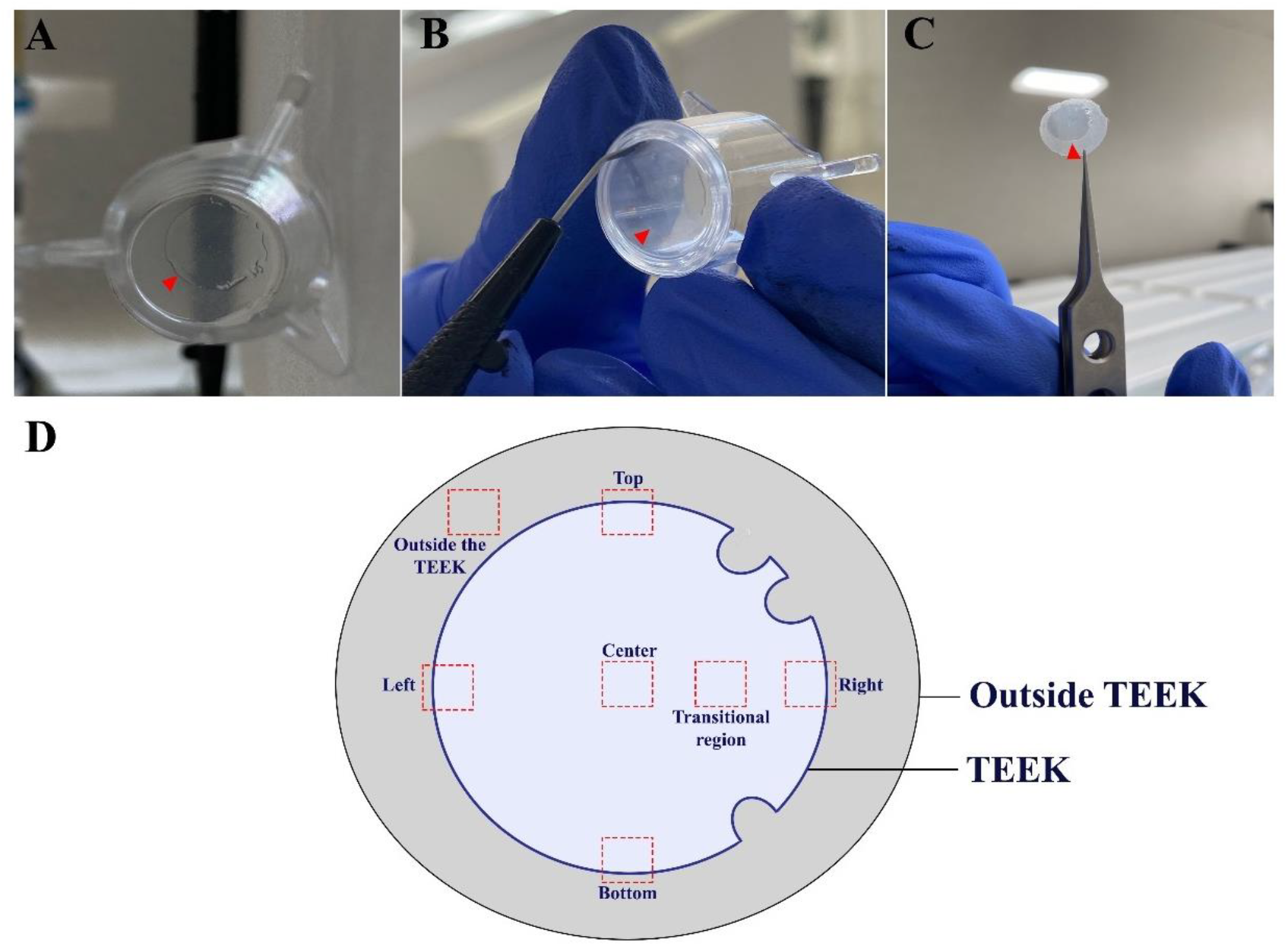
Tissue engineered endothelial keratoplasty (TEEK) after immunostaining. A. TEEK adherent to the bottom of an insert. B. Cutting of the insert membrane without touching the TEEK. C. Membrane and TEEK ready to be mounted between slide and coverslip. D. Diagram of the different observation zones on and outside the TEEK.

### Cellular segmentation and morphological analysis

As described recently in ^11^ we tuned a Cellpose model^12,13^ for segmenting NCAM-labelled hCECs. Cellpose is a generalist cell and nuclei segmentation algorithm with multiple pretrained model each trained on thousands of cells but which was not adapted to NCAM-labelled hCECs hence the need to tune a model specifically for this labelling on these cells which is largely facilitated by an easy to use graphical user interface. We developed a mathematical analysis python script (v3.11.4) to measure 5 parameters on segmented images: coefficient of variation of cell area (CV in %), adjusted CV (in %), percentage of cells with 6 neighbors (HEX), HEX-Q (%), filimorphism (%) to conduct extensive cell polymegathism and pleiomorphism analysis (Figure 3). We also used an Endothelial Quality Score (EQS) to evaluate the global morphology of TEEK hCECs. This scoring system assigned 50% weight to ECD, as it is the primary criterion used in clinical practice (eye banking), and 50% to morphology, considering that both were equally important. Morphology evaluation was based on the five parameters previously described: HEX, HEX-Q, CV, adjusted CV and filimorphism, each contributing 1/10 of the EQS. The EQS was derived from Z-scores, which were calculated by the following formula: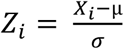 where *Z*_*i*_ was the score value, Xi the raw value, μ the population mean for the given criterion, and σ the standard deviation of the referred population. The higher the EQS, the better the endothelial quality.

**Figure 3.**
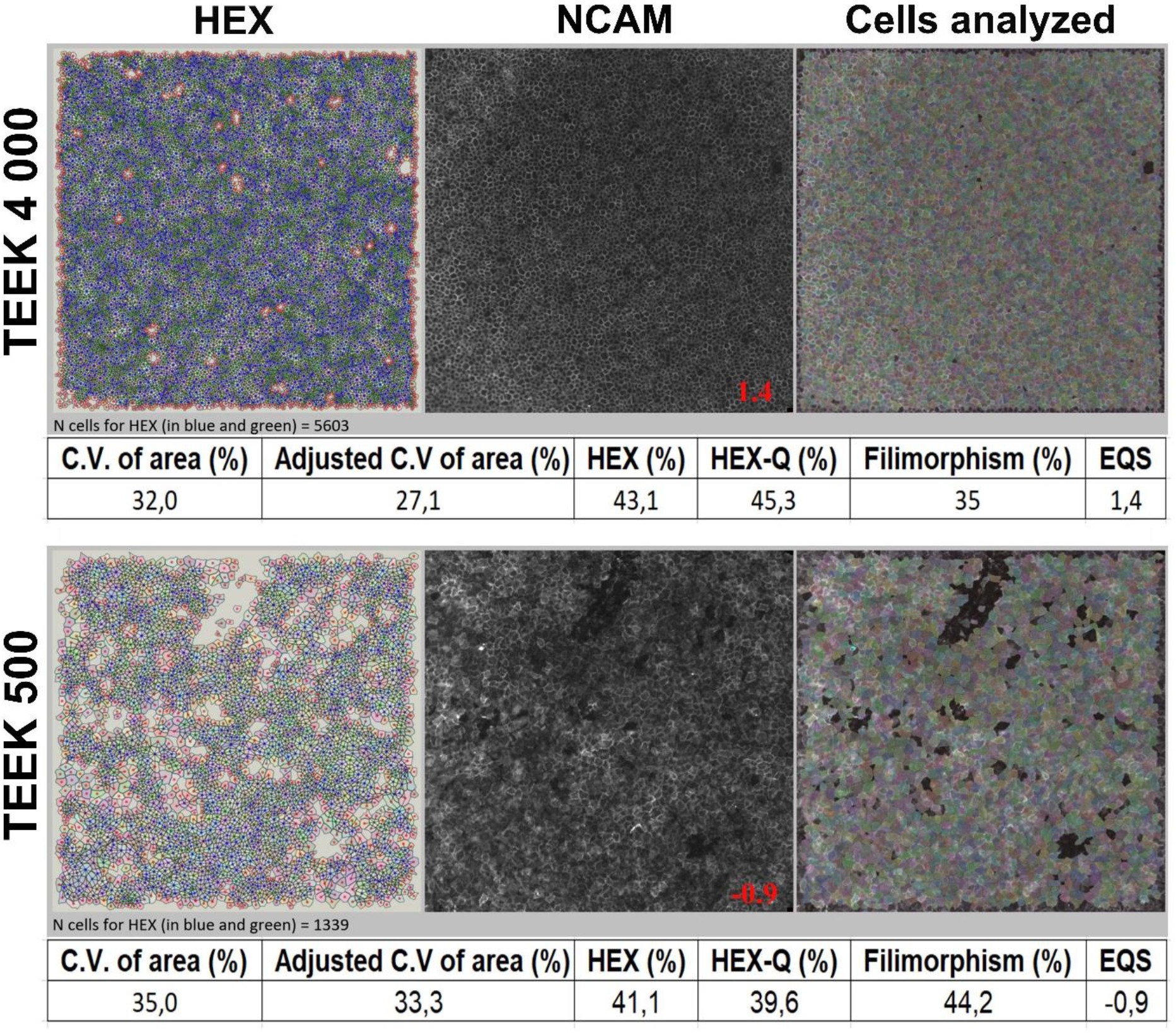
Example of Cellpose segmentation applied to NCAM labelled hCECs with computed morphometric parameters: CV, Adjusted C.V, HEX, HEX-Q and filimorphism (in %) and Endothelial Quality Score (EQS) for global morphology evaluation.

### Statistical analysis

Statistical analysis was performed using IBM SPSS Statistics 30.0.0.0. Given the sample sizes of the 3 groups, comparisons were carried out using the non-parametric Kruskal-Wallis test with Post-Hoc Dunn’s test using a Bonferroni corrected alpha for multiple comparisons (pairwise group comparisons) to evaluate the influence of the LCD side, as well as for the comparison of morphometric criteria, EQS, and vECD.

### Experiment

After 28 days of culture, for each of the 3 TEEK groups, morphological criteria, EQS and vECD did not differ significantly between the hCECs adhering to the LCD and those adhering to the membrane of the culture insert outside the TEEK. Similarly, for each of the 3 TEEK groups, vECD did not differ significantly between the 2 sides of the LCDs (data not shown). The data obtained on the anterior and posterior sides were therefore grouped for the subsequent comparisons between the 3 groups. The data were therefore obtained by analyzing 7739+/-2528 cells (min 3500 - max 11875) per TEEK. All TEEKs showed a continuous endothelium (Figure 4). As expected, the vECD of the TEEK 500 was significantly lower than that of the other two groups (P=0.005). None of the 5 morphometric parameters taken separately differed between the 3 groups, but the EQS score was significantly lower for the TEEK 500 than for the TEEK 4000 (P=0.0037). Between the TEEK 2500 and 4000, none of the 6 criteria differed significantly. All of these results were shown in (Figure 5).

**Figure 4.**
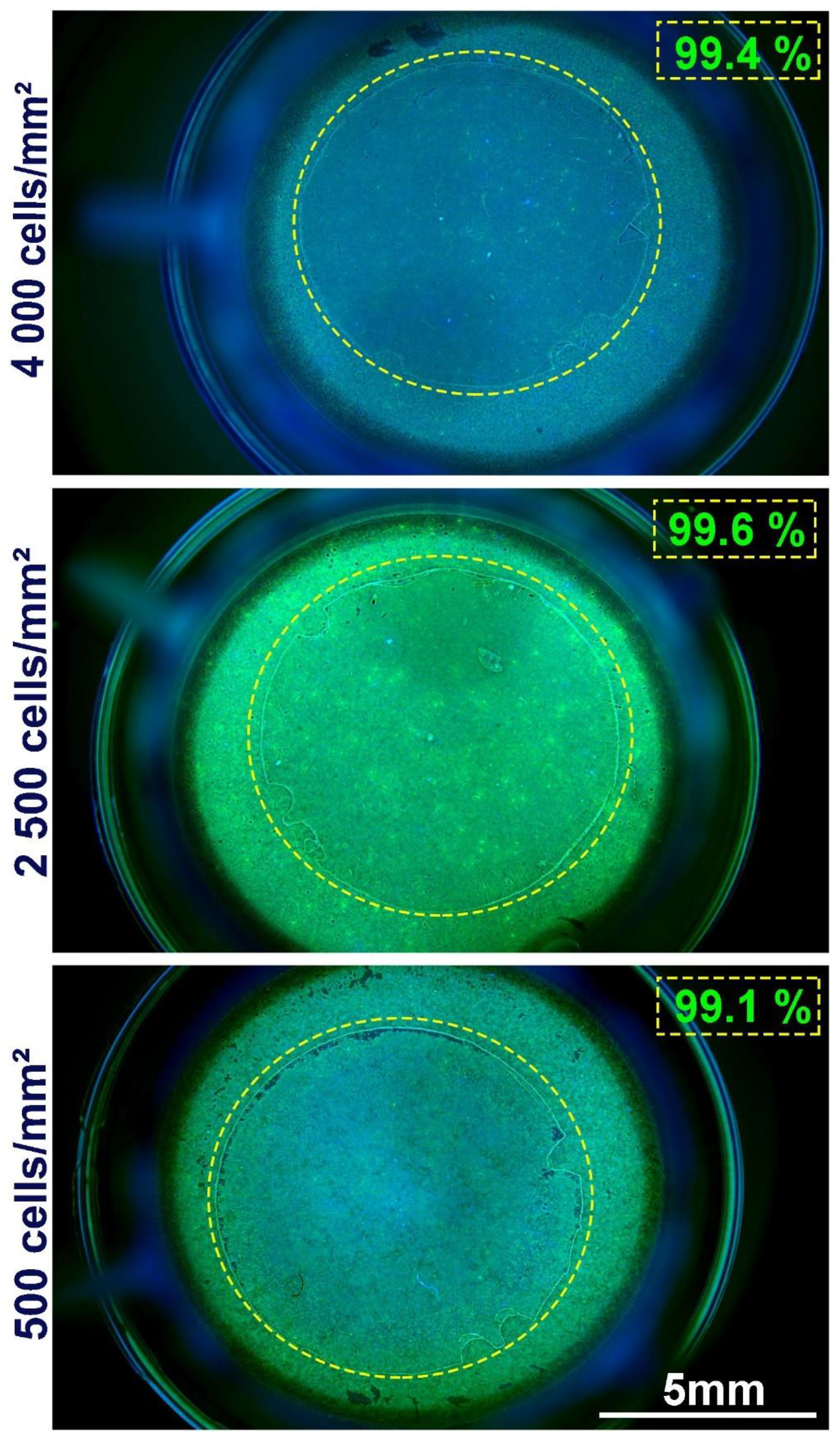
Uniformity and viability after 28 days of Tissue Engineered Endothelial Keratoplasty (TEEK) with 3 increasing seeding densities. Staining with Hoechst 33342 and Calcein-AM (green fluorescence) directly in their culture insert. Observations under macrozoom microscope (x1). Yellow dotted lines outlining the TEEK. Viability expressed as % of the TEEK surface.

**Figure 5.**
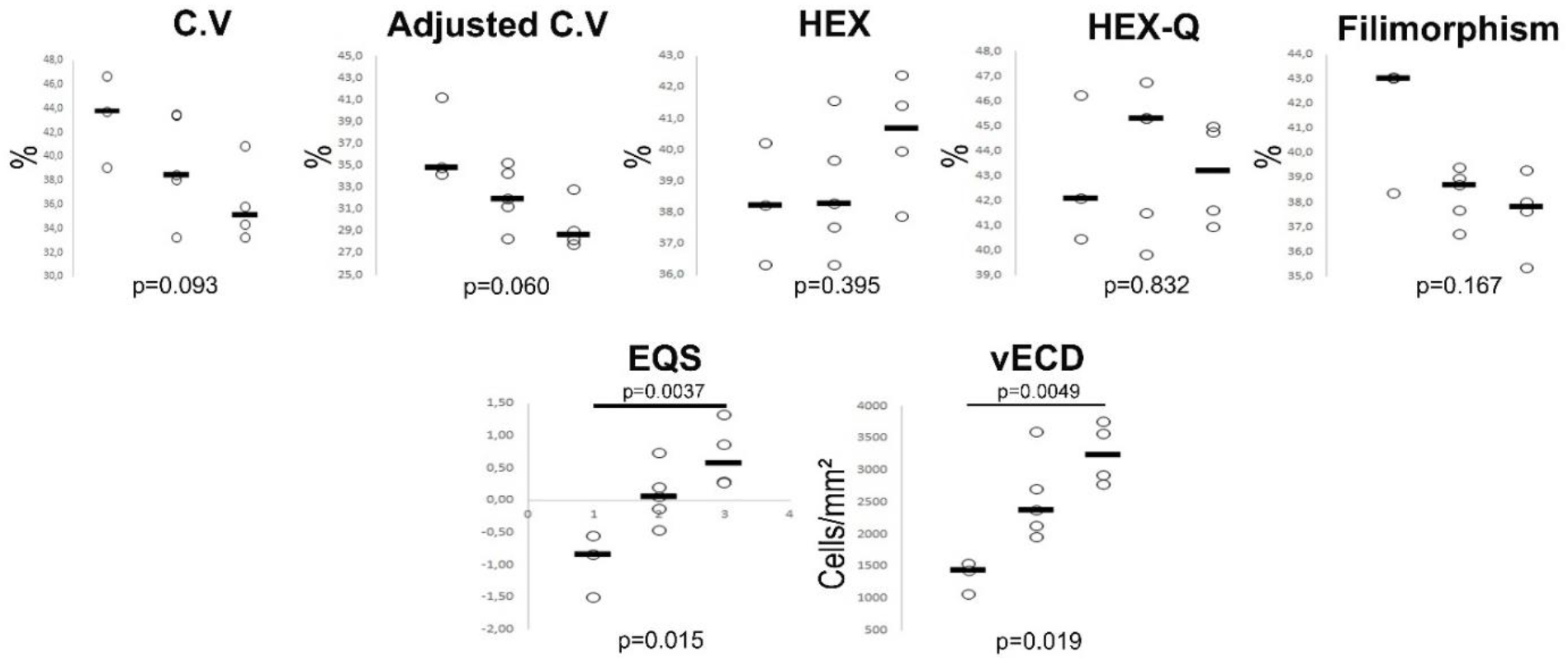
Comparison of endothelial morphometry and viable corneal endothelial cell density (vECD) between the 3 groups. Individual values and median (horizontal bar). C.V: coefficient of variation of cell area, HEX: hexagonality, HEX-Q: corrected hexagonality, EQS: Endothelial Quality Score.

## Discussion

In the strategy of replacing donor corneas with alternative solutions, TEEK faithfully replicates the Descemet Membrane Endothelial Keratoplasty (DMEK) endothelial graft. It consists of a biocompatible support, a LDC in our case^3,5^, on which a monolayer of hCEC adheres, forming a neo-endothelium capable of restoring the corneal histology in its entirety. In this pilot study, we demonstrate that it is possible to control the final endothelial cell density (ECD) during the bioengineering process and obtain grafts with supra-physiological ECD.

We used an original method of femtosecond laser capsulotomy that ensures the reproducible production of 8 mm diameter LCDs with minimal manual intervention and a conventional method of hCEC culture^5^. Note that we have obtained that the lens can now (since Nov 2024) be harvested for therapeutic purposes from deceased individuals ^14^.

For the final quality assessment of TEEK, we measured the vECD^8^, which stands as the best quantification of the truly useful cellular capital (37 referenced articles since our initial description in 2010^8,15^), and used an innovative method based on an AI image analysis algorithm to determine an endothelial quality score from a very large number of automatically segmented cells. The EQS can be considered as an automatic and objective assessment of endothelial mosaic quality, usable as a new biomarker, particularly for spotting hCECs in mesenchymal-to-endothelial transition. This new test has currently been trained on immunostaining images of NCAM (destructive test). During the production of clinical batches, it could be used on a sacrificed sample or needs to be adapted to images without staining or with a non-toxic stain to be used on each TEEK. It will complement the critical quality attributes (CQA) used to release cell batches ^16^.

This work helps to clarify several critical points of the bioengineering process. When hCECs are seeded at 500 cells/mm^2^, they reach a DCE close to their initial DCE before replanting on the LCDs after 28 days. However, it is possible to achieve a higher final DCE than before replanting by seeding a larger number of cells on the LCDs. This concentration effect allows for modulation of the final quality of the TEEK, but notably, there seems to be a maximum limit as none of the 4 TEEK 4000 exceeded 3800 viable cells/mm^2^. It is likely that this limit is related to the DCE before replanting (1865 cells/mm^2^ in this series), but this will need to be verified experimentally. Notably, the achievement of TEEK with a vECD >2000 cells/mm^2^ using donors aged 83 and 88 years is an important breakthrough, as up until now, clinical-grade cultures have been obtained from donors under 30 years of age^2 17^. Finally, we have clarified that the orientation of the LCD (anterior or posterior face) does not seem to influence the adhesion and growth of hCECs. To our knowledge, this result has not been known until now ^18,19,20^

TEEK presents several theoretical clinical advantages over the injection of cells in suspension: 1) the presence of a structured basement membrane on which hCECs reform a confluent endothelium; 2) the possibility of maintaining cultures to allow for the maturation of various intercellular junctions. These two characteristics should enable early endothelial function recovery similar to a current DMEK. Furthermore, TEEK transplantation allows for an exact replication of the current surgical procedure, which involves the removal of the patient’s pathological endothelium, whereas it is preserved in the injection therapy; 3) elimination of the risk of a large number of cells being carried into the general circulation (out of the 1 × 10^6^ injected cells, just under half reform an endothelium of 110 mm^2^ with a maximum of 4000 cells/mm^2^). Even though this situation has not caused any adverse effects in preclinical ^21^ and clinical ^2,17,16^ studies of suspension cell injection therapy, this notable difference should facilitate the approval of this ATMP by health authorities; 4) control over the number of adherent cells delivered to the patient without the need to inject a molecule promoting cell adhesion (such as Rho-Kinase inhibitors injected with suspension cells); 5) similarly to DMEK, TEEK should be transportable from the manufacturing site (tissue therapy unit) to the operating room under simple conditions, potentially pre-loaded and ready to be transplanted ^22^.

These various theoretical advantages will, of course, need to be verified by clinical data.

In clinical practice of endothelial grafts, at the request of surgeons, cornea banks reserve corneas with high DCE for this technique. While the validation threshold for a conventional full-thickness graft was set about 40 years ago at 2000 cells/mm^2^ (a pragmatic threshold balancing satisfactory clinical results and an economically acceptable proportion of donor corneas), and is nearly universal worldwide, it is most often set at 2400 cells/mm^2^ for endothelial grafts, as the preparation and surgical technique involve more manipulations that cause additional cell loss, which varies depending on the difficulties encountered on a case-by-case basis. Here, we show that TEEK bioengineering should allow achieving even higher DCE levels capable of competing with the postoperative DCE of 4000 cells/mm^2^ reported 2 years after treatment with suspension cell injection ^23^. This supra-physiological DCE should allow for better resistance to intra- and postoperative events, thereby prolonging graft survival, leading to the concept of the “super TEEK.” Moreover, the ability to customize TEEK production also opens the perspective of personalizing graft allocation to the recipient while optimizing the limited hCEC resource: for example, a TEEK of 1500 to 2000 cells/mm^2^ could most likely result in excellent visual recovery in an 85-year-old patient or older, as demonstrated by hemi ^24^ and quarter DMEK ^25^.

The main weakness of this pilot study is its limited power. In particular, it was not possible to differentiate between TEEK 2500 and 4000. The accumulation of data on a larger scale, for example, during an industrialization process, will allow for a multivariate analysis of the influence of various parameters concerning cell culture (donor age, DCE before culture, method and duration of preservation, composition of the culture medium, seeding density, and maturation time after seeding).

## Conclusion

The bioengineering of endothelial grafts on LCDs allows for the control of the number of viable cells after 4 weeks and, consequently, the creation of a range of TEEKs potentially enabling personalized allocation to the recipient. It also allows for the production of grafts with supra-physiological DCE. These “super-TEEKs” could have prolonged survival in recipients.

## Acknowledgments

We would like to thank the Fondation des Aveugles de Guerre (Paris) for their support (one-year PhD scholarship for Ms. Inès Aouimeur). We extend our sincere gratitude to the cornea donors and their families for their generous and invaluable contributions to medical and scientific advancement. We also wish to sincerely thank the Eye Banks of Saint-Étienne, Besançon, and Nantes for their crucial support in providing corneas for research purposes. We also thank the transplant coordination teams of the CHU of Saint-Étienne and Nantes, as well as the laboratory of anatomy at the Faculty of Medicine of Saint-Étienne.

## Authors contribution

Conception : Inès AOUIMEUR, Zhiguo HE ; Anaick MOISAN, Philippe GAIN, Gilles THURET, Corantin MAURIN

Experimentation : Inès AOUIMEUR, Louise PARVEAU, Sofiane FRAINE, Guillaume BONNET, Tomy SAGNIAL, Gauthier TRAVERS, Sédao XXX

Interpretation of Results: Inès AOUIMEUR, Zhiguo HE, Cyril MAUCLAIR, Gilles THURET, Corantin MAURIN

Drafting : Inès AOUIMEUR, Zhiguo HE, Philippe GAIN, Gilles THURET, Corantin MAURIN

Editing : Anaick MOISAN, Philippe GAIN, Gilles THURET, Corantin MAURIN Final approval : all authors

