## Supplemental data for "Tissue engineered endothelial keratoplasty with controlled endothelial cell density: proof of concept and paving the way for super TEEKs"

**Supplementary data**

**Table S1.** Characteristics of the lens capsules used

| **ID** | **Sexe** | **Age (y.o)** | **Death-to-retrieval time (hours** |
| --- | --- | --- | --- |
| 1 | M | 56 | 23 |
| 2 | M | 56 | 23 |
| 3 | M | 65 | 12 |
| 4 | M | 77 | 1 |
| 5 | M | 77 | 1 |
| 6 | F | 95 | 20 |
| 7 | F | 95 | 20 |
| 8 | F | 90 | 37 |
| 9 | M | 69 | 29 |
| 10 | M | 69 | 29 |
| 11 | M | 71 | 2 |
| 12 | M | 71 | 2 |
| 13 | F | 70 | 62 |
| 14 | M | 82 | 0 |
| 15 | M | 89 | 5 |

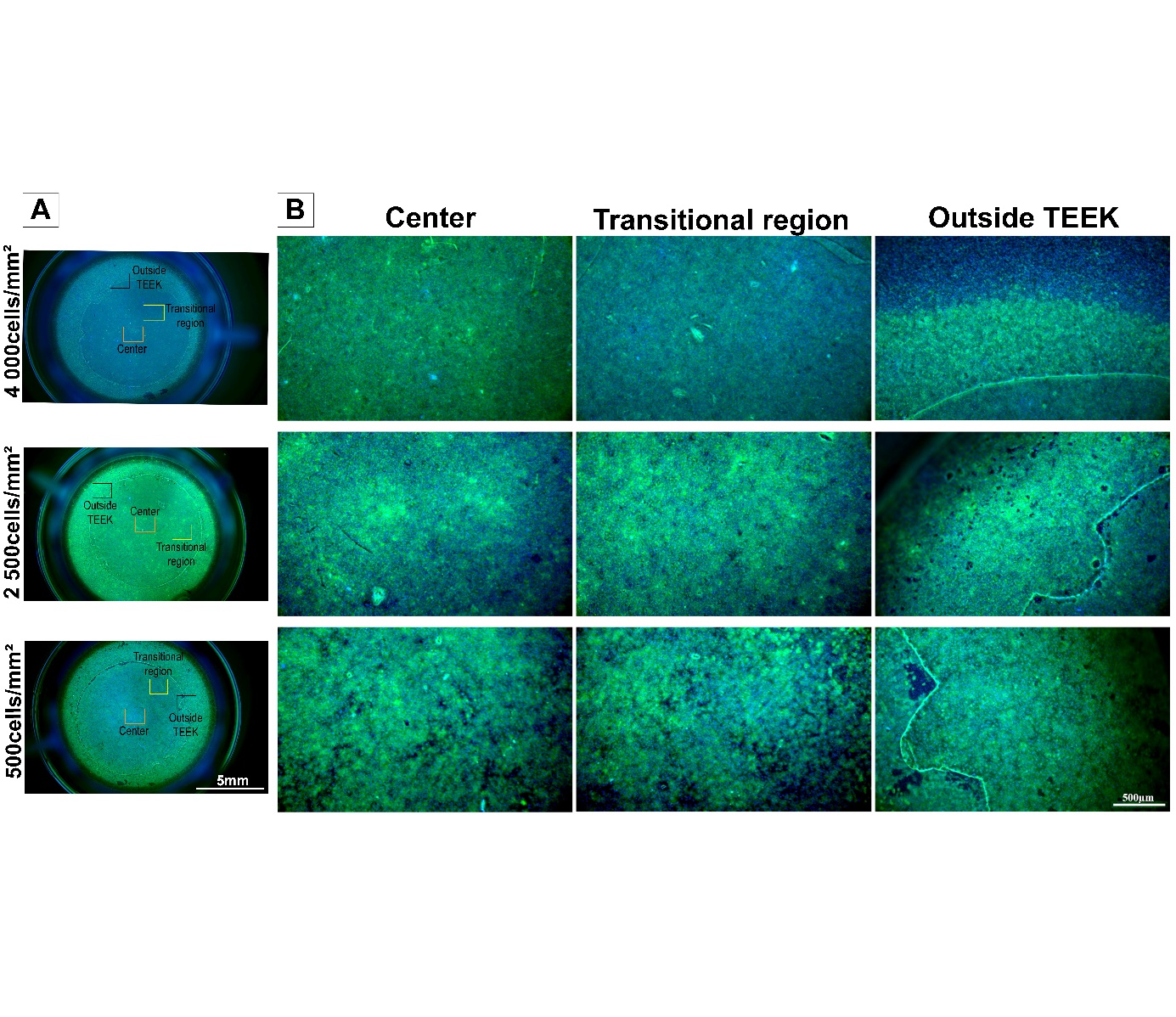

***Figure S1.*** **Microscopic Observation of Different Types of TEEK in Inserts** A. Observations of the entire inserts (at x1 magnification). B. Observations of the different areas inside and outside the TEEK (at x6.3 magnification).

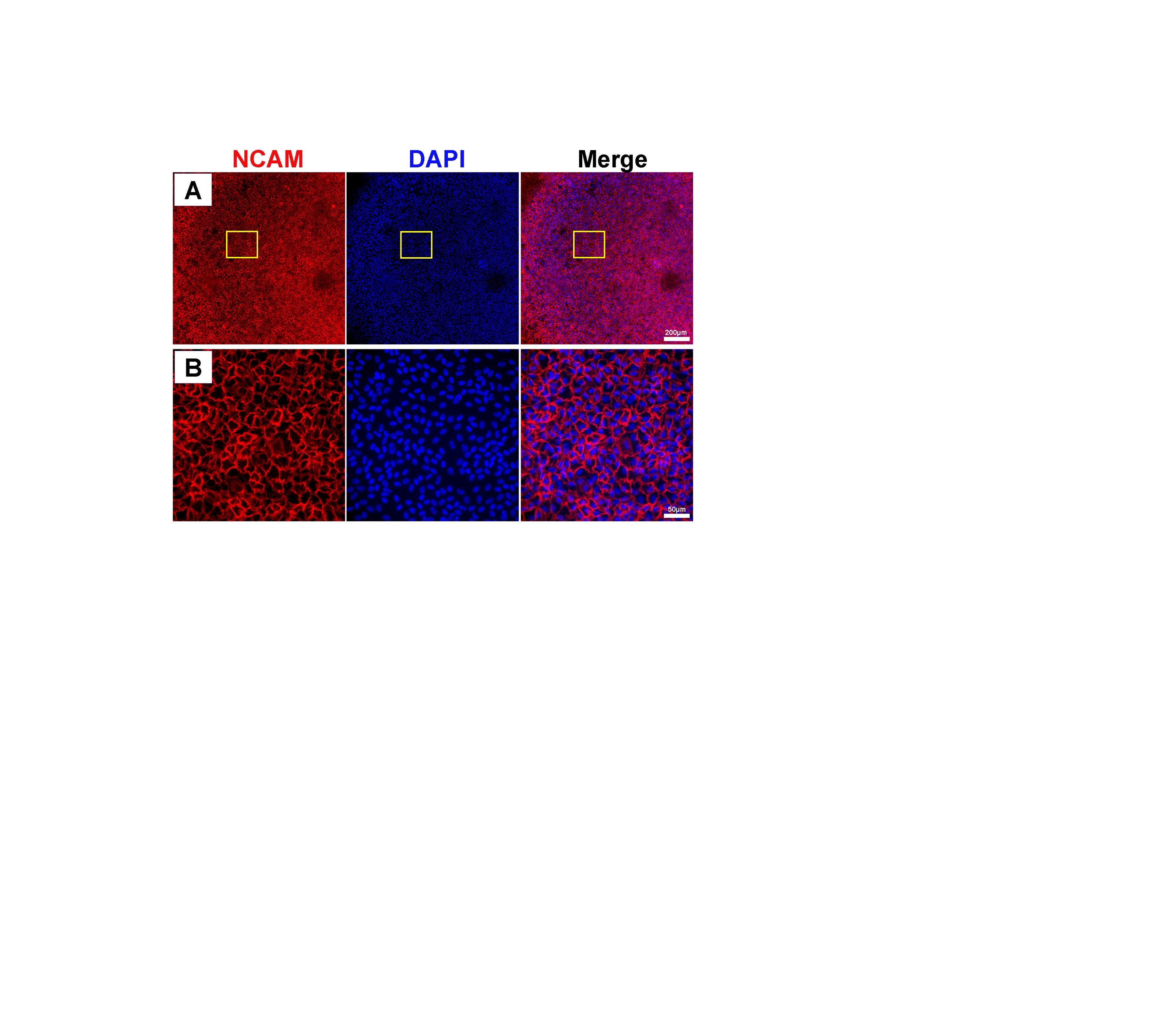

**Figure S2.** Observation of a TEEK at 4,000x under an Inverted Epifluorescence Microscope.In red, the NCAM specifically labels the cell membranes of hCECs, and in blue, DAPI stains the cell nuclei. A. Observation with a x10 objective. B. Observation with a x40 objective.
